# Molecular Fossils from Microorganisms Preserved in Glacial Ice

**DOI:** 10.1101/019240

**Authors:** P. Buford Price, J. Jeffrey Morris, Ryan C. Bay, Ajeeth Adhikari, Stephen J. Giovannoni, Kevin L. Vergin

## Abstract

The study of microbial evolution is hindered by the fact that microbial populations leave few fossils. We hypothesized that bacterial cells preserved in ancient ice could be used as a molecular fossil record if their DNA could be extracted and sequenced. Channels formed along triple junctions of ice crystals contain liquid “veins” in which microbial cells may be preserved intact. Since vertical motion through the ice matrix is impossible, microbes found in ice cores are representative of microbes present at the time the ice was formed. We detected chlorophyll fluorescence in intact ice cores taken from Greenland and Antarctica. Flow cytometric analysis localized at least some of this fluorescence to particles < 1 μm in diameter. Metagenomic analysis of meltwater indeed revealed sequences similar to modern strains of the picocyanobacterial genera *Synechococcus* and *Prochlorococcus*, and some of these sequences were distinct from any sequences known from modern oceans or glacial environments. Our study is a first proof-of-concept of the use of ice cores as records of microbial evolution, and we suggest that future genetic studies with higher vertical resolution in the cores might shed light on the pace and character of evolution of these ecologically important cells.

## INTRODUCTION

Studies of the evolutionary history of life on Earth are often informed by a combination of fossil evidence and molecular phylogenetics. The former method allows robust estimates of the timing of major evolutionary events, but usually is limited to hard-bodied organisms and, like all purely morphological studies, is prone to researcher bias. Molecular comparisons (e.g., of homologous DNA sequences) are much more amenable to statistical examination but are generally only available for extant organisms. “Molecular fossils” – e.g., ancient DNA that remains sufficiently intact for analysis – are rare, due to the continuing degradation of large biomolecules after an organism’s death. Most studies of ancient biomolecules either deal with a small number of individuals [e.g. studies of mammoth and Neanderthal DNA, 1, 2] or with mixed populations that are difficult to connect directly with any extant species [e.g. bacterial communities in Antarctic ice, 3]. It remains difficult to extrapolate from these studies to large-scale population genetic questions, particularly of biogeochemically important microbial populations.

Here we explore the feasibility of studying the long-term evolution of microorganisms by exploiting the discovery of chlorophyll-containing cells in ice cores from Antarctic and Arctic glaciers [4]. Microbial cells may become preserved in nutrient-rich liquid veins within the ice matrix at temperatures too low for either growth or significant movement in the ice matrix [5]. Therefore, cells in a given ice stratum are representative of strains that were alive in the oceans at the time of deposition onto glacial ice. The cell envelopes of a large fraction of the preserved cells remain intact [4], leading to the possibility that their genomes may also remain intact. The recovery of high-molecular weight DNA from ancient, datable microbial communities with readily identifiable modern relatives would constitute a fossil record of microbial evolution, with the oldest genomes at the bottom of an ice core and the youngest at the top.

In this manuscript we present results from our first attempt to extract and sequence microbial DNA from ice cores from Antarctica and Greenland. Despite substantial numbers of sequences from probably contaminants, we were nevertheless able to confirm the presence of unicellular marine cyanobacteria closely related to the widespread modern taxa *Synechococcus* and *Prochlorococcus* in 3 of the 4 ice cores we examined. We believe these results are a proof-of-concept of the use of ice cores as records of microbial evolution, and based on our estimates of cyanobacterial abundance throughout the depths of both Greenland and Antarctic ice and the time frames represented by those cores, we expect future, refined efforts to allow evaluation of the evolution of marine bacteria on a million-year timescale.

## RESULTS AND DISCUSSION

### Analysis of Chl fluorescence in ice cores

Using a scanning fluorometer (the BFS, see Methods), we detected Chl fluorescence throughout intact ice cores from the Western Antarctic Ice Sheet Divide Ice Core project (WDC) and the Greenland (GISP2D) project. Chl fluorescence in WDC ice sharply decreased in the top 200 m, followed by a relatively abrupt shallowing of the slope from 1200 m down to ∼3100 m. (Fig. 1A). Similar trends were observed in intact GISP2D cores (Fig. 1A).

**Figure 1.**
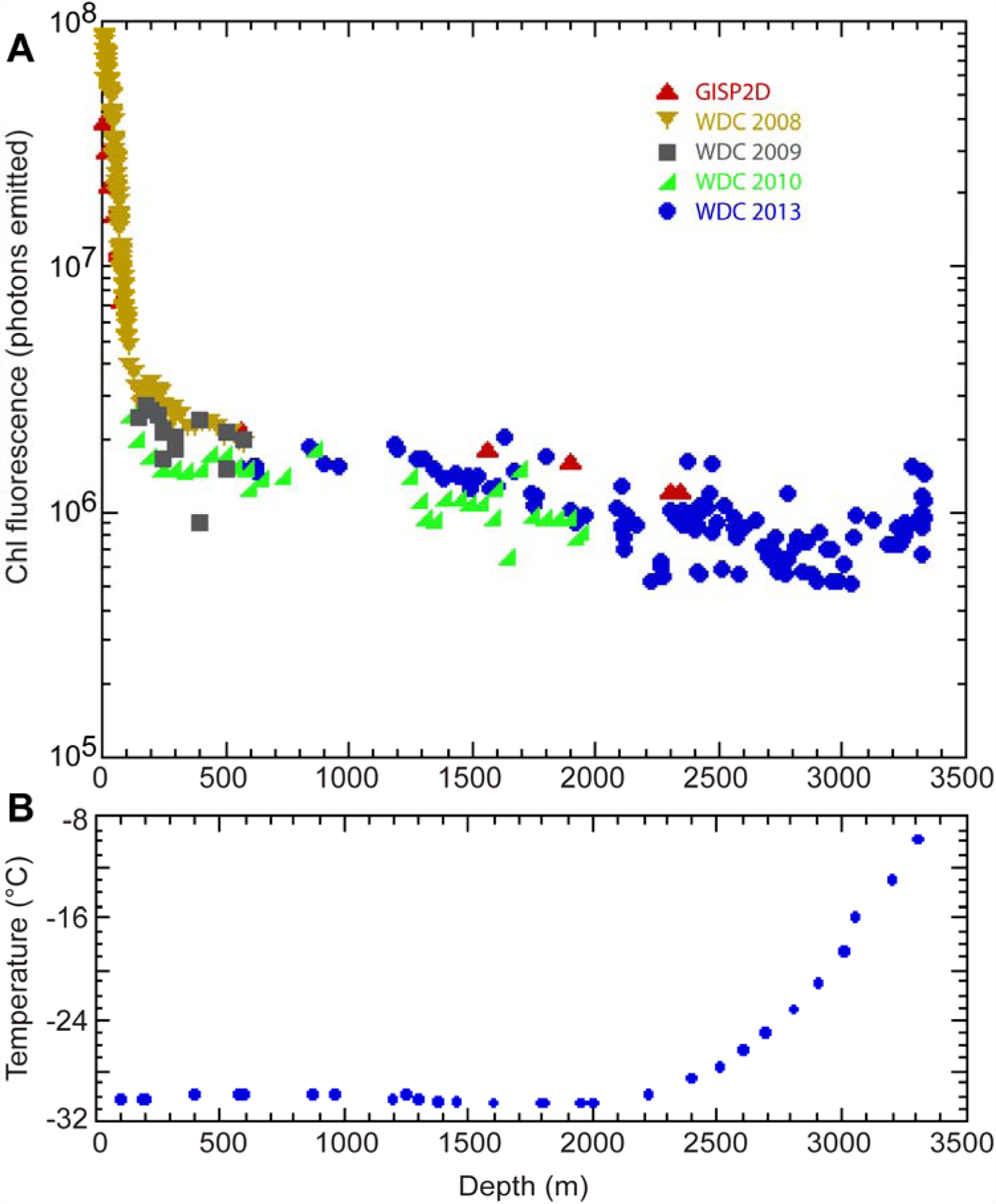
Fluorescence in intact ice cores. A) Normalized Chl autofluorescence (number of photons emitted between translations of the horizontal stage) for WDC ice examined at NICL in 2008, 2009, 2010, and 2013, and for GISP2D ice examined in 2008. B) Temperature in the same borehole from which the WDC core was removed.

FCM of melted ice from WDC, GISP2D, and numerous other cores (Table 1) revealed micron-sized Chl-containing particles similar in size and shape to marine cyanobacteria of the genera *Prochlorococcus* and *Synechococcus* in every sample that we analyzed (e.g. Fig. S1), suggesting that at least some of the Chl detected by the BFS was contained in intact cells. Events in the Chl/SSC gate calibrated with cultured marine picophytoplankton were extremely rare in sterile controls, and all ice core samples were well above the average control value of 9.9 counts mL^-1^ (Fig. 2C).

**Figure 2.**
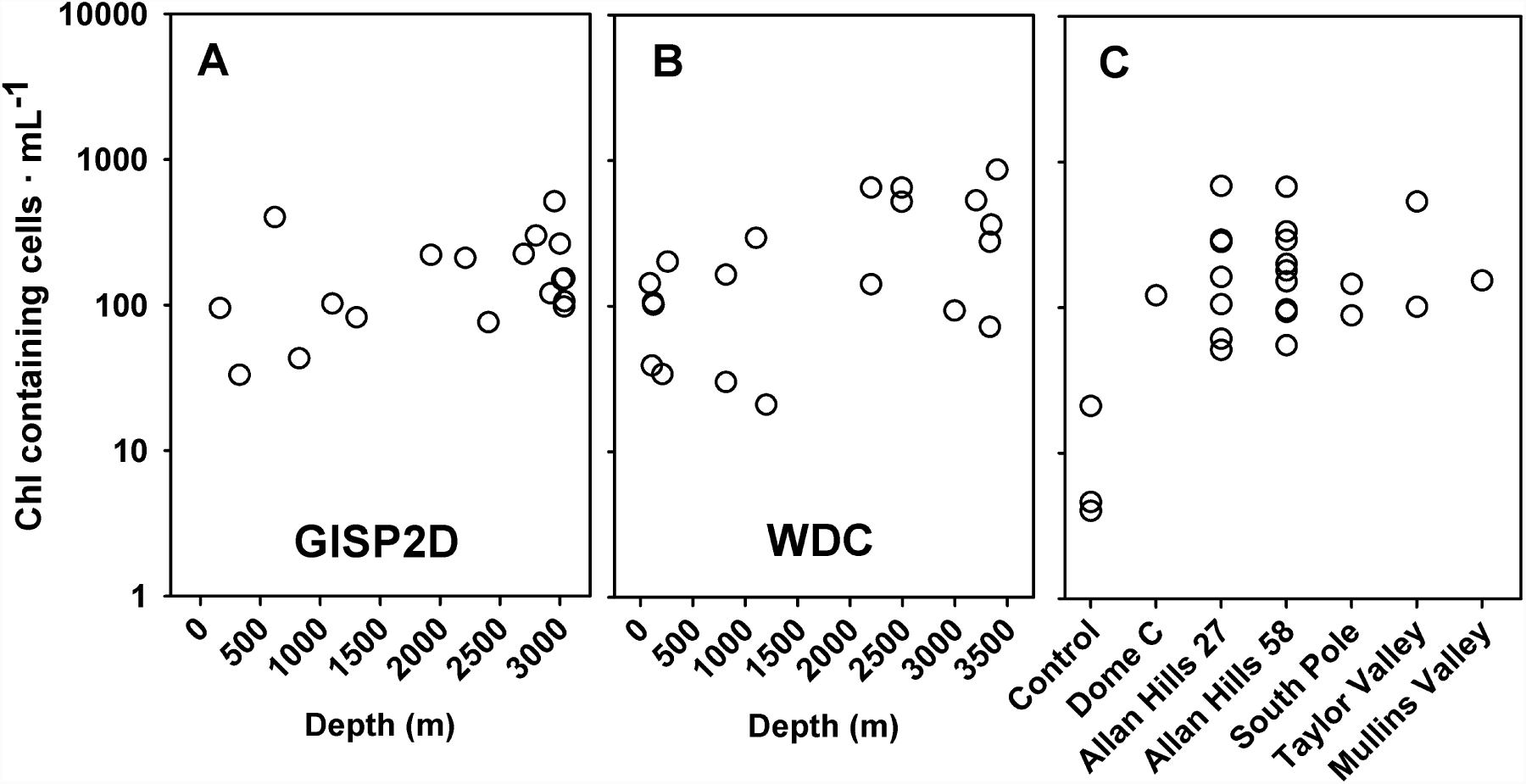
Cyanobacterial concentrations assessed by flow cytometry. Melted ice cores were passaged through a 1 μm filter prior to analysis. GISP2D (A) and WDC (B) ice were examined at many depths. Samples from additional cores (C) were examined at fewer depths (Table 1). “Control” in panel C refers to non-sterilized deionized water (2 lower points) and tap water (higher point) analyzed in the same manner as ice core meltwater.

**Table 1.** Ice samples studied.

| **Ice Core** | **Depths examined (m)** | **Age range (ya)** | **Source** | **Assays performed** |
| --- | --- | --- | --- | --- |
| Western Antarctic Ice Sheet Divide (WDC) | 86-3402 | 250-68,700 [36] | NICL | BFS, FCM, Genomic (265-269, 2487-2497 m) |
| Greenland Ice Sheet Project 2 (GISP2D) | 19-3037 | 200-162,600 [36] | NICL | BFS, FCM, Genomic (320-322, 1914-1915 m) |
| Dome C (Antarctica) | 120-188 | 3,100 [36] | E. Wolff | FCM |
| South Pole | 111-179 | 1,030-1,840 [37] | NICL | FCM |
| Vostok Station (Antarctica) | 144-214 | 5,500-9,000 [36] | NICL | EFM |
| Allan Hills (BIT58, Antarctic Blue Ice) | 26-177 | ~350,000 (ages uncertain) [38] | A. Kurbatov | FCM |
| Allan Hills (BIT27, Antarctic Blue Ice) | 105-289 | ~140,000 (ages uncertain) [38] | A. Kurbatov | FCM |
| Taylor Valley (Antarctic Dry Valley) | 28-440 | Unknown | J. Severinghaus | FCM |
| Mullins Valley | 9 | ~1,000,000<sup>1</sup> | D. Marchant | FCM |

<sup>1</sup> Michael Bender, personal communication; based on $^{40}\text{Ar}/^{38}\text{Ar}$ dating. BFS, Berkeley
Fluorescence Spectrometer; FCM, Flow Cytometry

In contrast to the data from intact cores (Fig. 1A), the abundance of Chl-containing particles increased with depth for both GISP2D (Fig. 2A, Spearman ρ = 0.332, P of true ρ < 0, 0.089) and WDC ice (Fig. 2B, ρ = 0.494, P of true ρ < 0, 0.013). The opposite trends in intact core Chl fluorescence and abundance of Chl-containing cells counted by FCM was possibly explained by a decrease in the fluorescence intensity of individual particles with depth for GISP2D ice (Fig. S2A, Spearman ρ = -0.448, P of true ρ > 0, 0.031); however this trend was not observed for WDC ice (Fig. S2B, ρ = 0, P of true ρ > 0, 0.5). Another explanation derives from the differential preservation of different size classes of chlorophyll-containing cells. Larger photoautotrophs such as diatoms and seaweed fragments would be excluded from ice veins and would likely die during ice crystal formation. Chl fluorescence from these non-viable cells would slowly decay, explaining the downward trend in whole-core fluorescence (Fig. 1), but importantly none of the Chl from these cells would be detected by our FCM methods. In contrast, some small cells, including the smallest phytoplankton, would essentially be cryopreserved within the veins and these cells would comprise a larger fraction of total Chl as the larger cells decayed.

### Phylogenetic placement of microorganisms preserved in polar ice

We successfully amplified DNA from all four depths (two from the GISP2D and two from the WDC ice cores) that we attempted (Table 1). A total of 55,505 quality-controlled pyrosequences were examined (Table S2), of which nearly 50% belonged to NTUs [nodal taxonomic units, 6] that were present in negative control samples and were thus removed. Of the remaining 28,336 sequences, nearly 97% belonged to NTUs that were closely related to *Escherichia coli*. Few *E. coli-*like sequences were present in negative controls, suggesting that *E. coli* was present in or on the ice cores and was not effectively removed by our decontamination procedures. Due to their high abundance in the sequencing run relative to all other sequences as well as the ubiquity of *E. coli* DNA in biology labs, we reasoned that these *E. coli* sequences were almost certainly human-introduced contaminants and removed them from subsequent analyses. This level of contamination was unfortunate, but is not uncommon for studies of ancient DNA or other systems for which the yield of target DNA is low [7, 8].

The remaining 883 sequences were distributed among 16 bacterial taxonomic groups (Table S3). Taxonomic composition differed between the samples (Fig. 3A). Bacteria often associated with soil and freshwater (e.g. Actinobacteria, *Caulobacter*, TM7, Firmicutes) were numerically dominant in WDC samples and were also abundant in GISP2D samples (Fig. 3A). Sequences identified as *Prochlorococcus*, a highly abundant unicellular cyanobacterium from temperate and tropical waters, were discovered in three of four ice core samples, including both depths from GISP2D ice. Indeed, *Prochlorococcus*-like sequences were numerically dominant amongst the retained sequences from deep GISP2D ice (Fig. 3A).

**Figure 3.**
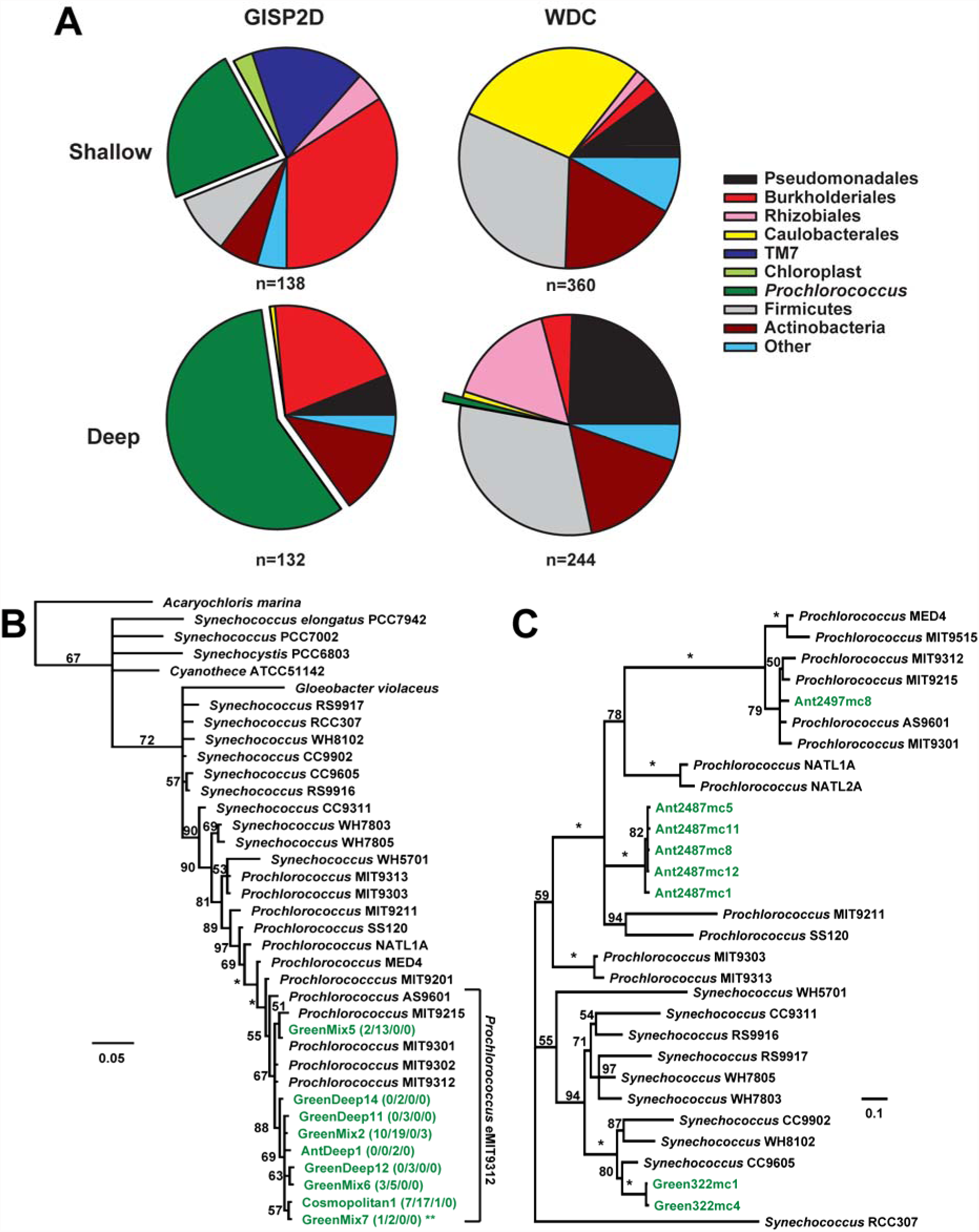
Phylogenetic analysis of cells recovered from ice cores. A) Phylogenetic placement of 16S pyrotag sequences from shallow and deep samples from GISP2D and WDC ice cores. n, number of quality sequences examined. B) Bayesian tree showing the relationship of the *Prochlorococcus* pyrotag sequences to other cyanobacteria. The tree was rooted on *Escherichia coli* MG1655 (branch pruned). Names in green are ice core sequences; the numbers that follow show how many times the sequence was observed in the different libraries (GISP2D Shallow/GISP2D Deep/WDC Deep/Unassigned). The double-asterisk by GreenMix7 indicates that this sequence had no exact match in any existing database. C) Bayesian tree showing the placement of ITS sequences from ice cores with other members of the *Synechococcus*/*Prochlorococcus* clade. For both trees, nodal values indicate posterior probabilities, expressed as percentages; * indicates 100% posterior probability. Scale bars indicate substitutions • bp^-1^.

Reasoning that marine cyanobacteria were unlikely to be contaminants, and since we saw evidence of *Prochlorococcus*-like cells by FCM, we looked more closely at these sequences. Placement of the 9 unique *Prochlorococcus*-like partial 16S rRNA pyrosequences in a phylogenetic tree (Fig. 3B) clearly placed these sequences within the eMIT9312 ecotype of *Prochlorococcus*, with an average genetic distance of 0.02 ± 0.005 (standard deviation) substitutions • bp^-1^ in comparison to the type strain MIT9312 (Table S4). Only one of the sequences had an exact match in a cultured strain (*Prochlorococcus* MIT9301), although 7 others had an exact match with uncultured bacteria from environmental studies (Table 2). One sequence, however, had no exact match in any of the databases we examined (Table 2). This sequence (marked with an asterisk in Fig. 3B) differed by a SNP and a single-base pair deletion from its closest match in the NCBI database. Moreover, it was detected independently in both GISP2D libraries, suggesting that these changes were not PCR or sequencing artifacts. Additionally, none of these sequences matched cultivated strains in use by our laboratories, nor were any *Prochlorococcus*-like sequences found in the 19,934 sequences found in negative controls, strongly suggesting that these warm-water cyanobacteria were in fact preserved within the ice cores.

**Table 2.** Best matches to ice core 16S rRNA and ITS sequences in various published databases. Values are % identical matches of best hit for the given sequence in the given database. Sequence names are based on the samples where the sequences were discovered and correspond to the names used in Fig. 3.

REFERENCES
| Sequence | nr | GOS | HOT | BATS | Bact | AntAq | IceMeta |
| --- | --- | --- | --- | --- | --- | --- | --- |
| **16S Sequences** |  |  |  |  |  |  |  |
| GreenDeep11 | 100 | 100 | 100 | 100 | 98.4 | 99.3 | 88.1 |
| Cosmopolitan1 | 100 | 99.5 | 99.8 | 99.5 | 97.9 | 98.8 | 87.8 |
| GreenDeep12 | 100 | 99.8 | 99.8 | 99.8 | 98.1 | 99.1 | 88.1 |
| GreenMix2 | 100 | 100 | 100 | 99.8 | 98.1 | 99.1 | 87.8 |
| GreenMix5 | 100 | 100 | 100 | 99.8 | 98.1 | 99.1 | 87.8 |
| GreenMix6 | 100 | 99.8 | 99.8 | 99.5 | 97.9 | 98.8 | 87.8 |
| AntDeep1 | 100 | 99.8 | 99.8 | 99.5 | 97.9 | 98.8 | 87.5 |
| GreenMix7 | 99.8 | 99.5 | 99.5 | 99.3 | 97.7 | 98.6 | 87.8 |
| GreenDeep14 | 100 | 100 | 100 | 100 | 98.4 | 99.3 | 88.1 |
| **ITS Sequences** |  |  |  |  |  |  |  |
| Ant2497mc8 | 99.9 | 99.1 | 98.3 | 97.3 | 97.1 | 97.8 | 85.3 |
| Grnld322mc4 | 96.3 | 94.3 | 91.9 | 92.5 | 91.6 | 88.2 | 86.7 |
| Grnld322mc1 | 96.8 | 96.2 | 92.1 | 92.2 | 92 | 87.9 | 86.4 |
| Ant2487mc5 | 90.7 | 95.2 | 88.4 | 87.1 | 95 | 95.5 | 86.8 |
| Ant2487mc11 | 89.9 | 92 | 82.5 | 87.6 | 90.9 | 93.1 | 86.9 |
| Ant2487mc8 | 89.4 | 95.9 | 82.5 | 87.5 | 90.8 | 92.8 | 86.9 |
| Ant2487mc1 | 88.4 | 81.9 | 80.3 | 85 | 89.4 | 91.3 | 93.4 |
| Ant2487mc12 | 90.5 | 85.9 | 84 | 84.9 | 86.8 | 85.4 | 89.1 |

We further recovered eight unique cyanobacteria-like ITS sequences from shallow GISP2D ice and deep WDC ice. Phylogenetic placement of these sequences (Fig. 3C) revealed greater diversity than the 16S sequences. The two ITS sequences from GISP2D clearly clustered with *Synechococcus*, a sister group to *Prochlorococcus*. One WDC sequence clustered in the same eMIT9312 *Prochlorococcus* ecotype as the 16S sequences. However, the other five WDC sequences formed a distinct clade within *Prochlorococcus* that was quite divergent from known sequences. While this clade was separated from all other sequences with 100% posterior probability, its branching order in relation to the other ecotypes of *Prochlorococcus* was unclear. Interestingly, three of these five sequences had better hits in an Antarctic metagenome [9] than in any other database (Table 2) suggesting the exciting possibility that currently undescribed cold-adapted picocyanobacterial strains might exist in these frigid regions.

### A fossil record preserved in glacial ice?

To glaciologists it might seem that cyanobacterial genomes would evolve little during only the last million years relative to their origin some 2.5 to 3 billion years ago. However, microorganisms are capable of very rapid evolution due to their vast population sizes and ability to undergo horizontal gene transfer during encounters with other cells in the ocean [10-13]. In contrast, growth, and hence evolution, effectively ceases after preservation in glacial ice. Also, the vertical distances bacteria may travel in veins is less than 1 μm yr^-1^, or < 10 cm in 100,000 yr [14]. Thus, microbial genomes in ice cores exist as molecular fossils, and their existence potentially opens a window into the microbial biogeochemistry of the past.

The rapid evolvability of bacterial populations is clear from evolutionary theory as well as from laboratory experiments [15], but it is difficult to observe it in action in situ. The time course and pace of microbial evolution have been documented in microbial pathogens [16], acid mine drainage communities [17], and laboratory populations [18]. The existence of a robust molecular fossil record linked to a well-studied modern microbial group would allow us to study these same processes on a much grander scale. *Prochlorococcus* and *Synechococcus* are globally distributed in oceans, and their modern population genetics are perhaps better understood than those of any other microbe in the environment. If our DNA sequences came from tropical cyanobacterial populations that were originally swept up from ocean surfaces by wind currents and deposited on ice thousands of miles away [4], then deeper examination of the microbiota of these ice cores should allow us to extend the ecological analysis of *Prochlorococcus* and *Synechococcus* to a multi-millennial or even million-year time scale. If instead we have discovered a novel clade of psychrophilic *Prochlorococcus*-like cyanobacteria, we can still study the pace of long-term divergence from, or introgression into, open ocean populations. Most importantly, because *Prochlorococcus* and *Synechococcus* are crucial players in Earth’s carbon cycle, their genome dynamics over successive glaciations might help us understand not only how microbes adapt to their changing environments, but also how the oceans will respond to ongoing anthropogenic changes. Our study represents a first step toward this ultimate goal, and we are optimistic that methodological refinements will greatly improve the resolution and reliability of future attempts to probe this valuable resource.

## METHODS

### Ice source and handling prior to analysis

We analyzed polar ice samples from a wide range of depths from the Greenland Ice Sheet Project 2 (GISP2D) and the Western Antarctic Ice Sheet Divide Ice Core (WDC) projects, as well as samples from the South Pole, Vostok Station, and a number of other sites in Antarctica and Greenland (Table 1). Fluorimetry analysis was conducted only on GISP2D and WDC ice, and was performed at the storage facility at the National Ice Core Laboratory (NICL) in Denver, CO. Small pieces of cores for use in flow cytometry and DNA analysis were stored in Price’s laboratory at −40° C until analysis.

### Non-destructive analysis of intact ice cores using the Berkeley Fluorescence Spectrometer (BFS)

Bramall [19] developed a scanning spectrofluorimeter powered by a 375 nm laser to map the 465 nm emission from the F420 pigment in methanogens inside solid ice cores at NICL [20]. During one week almost every year from 2008-2013, we were allowed to continuously occupy a room at -25°C at NICL that we were able to darken completely. One version of the BFS using a 7-channel 224 nm laser allowed tryptophan, chlorophyll (Chl), and volcanic ash to be mapped as a function of depth in ice cores. A strong peak in channel 6 was used to detect Chl and its decomposition product pheophytin [21] and strong emission in channel 7 was used to detect fluorescence from mineral grains transported from volcanic eruptions. With a laser beam ∼1 mm in diameter and brief bursts of laser light at intervals ∼1 mm perpendicular to the flat surface of an ice core cut longitudinally in half and polished, Price and co-workers were able to acquire 7 channels of data along a core ∼1 meter long in ∼2 minutes. Data from different scanning sessions were normalized as described previously [4, 20].

### Flow cytometry

Abundance of ice core bacteria were assessed using flow cytometry (FCM). In order to reduce surface contamination due to microbes not originating in the ice, we removed all but ∼4 ml from ∼50 ml ice samples with either stirred 6% household bleach or stirred sterile water. Then the remaining ice was melted and passed through a 1 μm pore size polycarbonate prefilter to eliminate volcanic ash particles, most mineral grains, and cells or cell debris too large to occupy veins in ice crystals. Microorganisms small enough to pass through this filter were analyzed with a Fortessa digital flow cytometer (BD). Chl autofluorescence was detected by emission at 670 nm (deep red) following excitation by a 488 nm argon laser, and phycoerythrin-like (PE) autofluorescence was detected by emission at 593 nm (orange) following excitation by a 567 nm laser. Particles were characterized in pairwise plots of Chl vs. PE and of Chl vs. side scatter (SSC, a measure of size and internal structure; see Fig. S1 for representative examples). Cyanobacterial gate boundaries were set using fluorescent beads of sizes 0.3, 0.5, and 1 μm. Cultures of *Prochlorococcus* (SS120, NATL1A, MIT9313, and MED4), *Synechococcus* WH8102, and *Ostreococcus* OTH95 and 601 were run as standards when initially adjusting voltage settings on the flow cytometer. After trial and error, events were triggered on values of SSC greater than 30 units, which accepted *Prochlorococcus* and *Synechococcus* cells and eliminated points labeled as noise (Fig. S1). FCM data were plotted and analyzed using Flowjo (TreeStar, Inc.).

Statistical analysis of FCM data was performed in the statistical computing environment R v. 2.15.1.

### DNA Extraction and Sequencing

In order to reduce the probability of contamination by foreign DNA on the exterior of the ice samples or on laboratory surfaces, all materials that came in contact with the ice core or melt water were sterilized either by heating at 450° C for 4 hours or by treatment with 10% HCl. Ice melting operations and meltwater filtering were conducted sequentially in a laminar flow hood. During the day of a run, several cores were removed from plastic bags, cut into samples of working volume ∼50 ml and rinsed with warm nanopure water to remove up to 50% of the volume of the outer surface. The remaining interior portion of the core was allowed to melt slowly in the dark. Microorganisms in the meltwater were collected on 0.1 μm polycarbonate membrane filters. Nucleic acids were extracted and separated as described elsewhere [22].

Two different amplification strategies were employed for DNA sequence analysis. Total community composition was assessed by amplifying partial 16S rRNA genes. These amplicons were processed as described elsewhere [6] except that a different reverse primer (519R appended with the 454B sequence (GCCTTGCCAGCCCGCTCAGTGWATTACCGCGGCKGCTG) was used. Pyrosequences were generated using Roche 454 GS FLX Titanium technology. Different barcoded forward primers were used for each sample, two nanopure controls, and a PCR negative control.

In order to target picocyanobacteria, the 16S/23S internal transcribed spacer (ITS) region was amplified using primers described previously (16S-1247F and 23S-1608R) that target *Synechococcus* and *Prochlorococcus* [23]. Samples with positive PCR signal were cloned into the pGEM-T-easy vector according to manufacturer’s instructions (Promega) and transformed into *E. coli* using an electroporator (Gibco). Purified plasmids from colonies containing inserts were screened by PCR using the universal internal priming sites 16S-1492F and 23S-241R and Sanger sequenced with these same primers using an ABI 373 sequencer.

Other studies have reported substantial degradation of DNA recovered from ancient ice [3, 24]. Our methods avoided this problem, however, by using an initial PCR amplification step, such that only reasonably intact DNA molecules were available for sequencing.

All sequence data have been deposited at Genbank as BioProject PRJNA239379.

### Phylogenetic analysis

16S pyrosequences were processed using the PhyloAssigner pipeline and reference phylogenetic tree described by Vergin et al. [6]. The taxonomic identification of the pyrosequencing reads (‘tags’) followed the approach of Sogin et al. [25]. The tags were compared by BLASTN [26] with a reference database of hypervariable region tags (RefHVR_V6, http://vamps.mbl.edu/) based on the SILVA database version 95 [27], and the 100 best matches were aligned to the tag sequences using MUSCLE [28]. Reference sequences were defined as those having the minimum global distance (number of insertions, deletions and mismatches divided by the length of the tag) to the tag sequence, and all reads showing the best match to the same reference V6 tag were grouped together as the same operational taxonomic unit (OTU) [‘best match definition’, 25, 29, 30]. Taxonomy was assigned to each reference sequence with the RDP Classifier [31]. This pipeline was, however, not precise enough to classify all bacteria from the deep Arctic Ocean, for which few sequence records are available in the databases [6]. To increase the taxonomic resolution, pyrosequencing reads were compared against our nearly full-length sequences using BLASTN. This additional step improved the classification up to the class/order level for 30,760 sequences (9.8% of the sequences) that were originally classified at the domain level, and 55,550 sequences (17.7%) that were first defined at the phylum level.

ITS amplicons from ice cores and metagenomes were aligned with a sample set of *Prochlorococcus* and *Synechococcus* ITS regions from fully-sequenced strains using MUSCLE. ITS alignments were then trimmed to remove 16S and 23S rRNA sequences as well as the two tRNA coding sequences [23]. Picocyanobacterial sequences identified from the pyrosequencing runs were aligned with *Prochlorococcus* and *Synechococcus* 16S sequences obtained from fully sequenced genomes using SINA [32]. Based on these alignments, we removed 16S pyrosequences that were potentially artifacts of the 454 sequencing process. Specifically, we eliminated any pyrosequences containing single nucleotide insertions relative to other *Prochlorococcus* or *Synechococcus* strains. We also eliminated any sequences that were only detected once. After this curation, 9 unique 16S pyrosequences and 8 ITS sequences remained for phylogenetic analysis.

16S and ITS alignments were used to infer phylogenetic relationships using a Markov chain Monte Carlo simulation as implemented in a multi-core build of MrBayes v3.2.1 [33]. Two parallel runs were used, each consisting of 1 “cold” chain and 3 “heated” chains. Analyses were carried out until the standard deviation between the runs was < 0.01. Phylogenetic trees were visualized using Geneious 5.5.9 (Biomatters).

BLAST analysis was employed to detect best matches for each of the ice core putative *Prochlorococcus*/*Synechococcus* 16S and ITS sequences remaining following curation. Sequences were compared against the NCBI nr database as well as a number of metagenomic datasets (Table S1). The raw read data from each metagenome was downloaded from CAMERA [34] and made into a BLAST database using BLAST+ v2.2.28 [35]. Ice core sequences were then used as queries against these databases using BLASTN and the single best match was returned. Because the average read length for most of the metagenomes was shorter than the sequences we were analyzing (Table S1), we split the sequences into sections of ∼240 bp and analyzed each section separately, then summarized the best hits for each fragment to get a total percent identity. Summarized BLAST results, alignment files, and tree files are available for download at datadryad.org (DOI Pending).

## ACKNOWLEDGEMENTS

We are grateful to the members of the WDC and GISP2 project members for their years of work collecting and preserving ice cores; Andrei Kurbatov, Jeffrey Severinghaus, David Marchant, and Geoffrey Hargreaves (curator, NICL) for providing ice samples; Richard E. Lenski for helpful discussions about evolutionary theory and population genetics; Riccardo Cavicchioli for informing us of the paper by Wilkins et al. (2013) on metagenomics of *Prochlorococcus* and *Synechococcus* in the Southern Ocean; Ben Temperton for assistance with 454 sequencing and the PhyloAssigner pipeline; and Kartoosh Heydari for assistance with the Li ka-Shing Fortessa flow cytometer. Price and Bay acknowledge partial support from NSF Grant ANT-1142178; Giovannoni and Vergin acknowledge support by a grant from the Marine Microbiology Initiative of the Gordon and Betty Moore Foundation to Giovannoni; Morris was partially supported by NSF Grants DBI-0939454 and OCE-1316101, and by a postdoctoral fellowship from the NASA Astrobiology Institute.

## References

1. Green RE, Krause J, Briggs AW, Maricic T, Stenzel U, et al. (2010) A draft sequence of the Neandertal genome. Science 328: 710–722.

2. Miller W, Drautz DI, Ratan A, Pusey B, Qi J, et al. (2008) Sequencing the nuclear genome of the extinct woolly mammoth. Nature 456: 387–U351.

3. Bidle KD, Lee S, Marchant DR, Falkowski PG (2007) Fossil genes and microbes in the oldest ice on Earth. Proc Natl Acad Sci USA 104: 13455–13460.

4. Price PB, Bay RC (2012) Marine bacteria in deep Arctic and Antarctic ice cores: a proxy for evolution in oceans over 300 million generations. Biogeosciences 9: 3799–3815.

5. Price PB, Sowers T (2004) Temperature dependence of metabolic rates for microbial growth, maintenance, and survival. Proc Natl Acad Sci USA 101: 4631–4636.

6. Vergin KL, Beszteri B, Monier A, Thrash JC, Temperton B, et al. (2013) High-resolution SAR11 ecotype dynamics at the Bermuda Atlantic Time-series Study site by phylogenetic placement of pyrosequences. ISME J 7: 1322–1332.

7. Salter SJ, Cox MJ, Turek EM, Calus ST, Cookson WO, et al. (2014) Reagent and laboratory contamination can critically impact sequence-based microbiome analyses. BMC Biology 12.

8. Shapiro B, Hofreiter M (2014) A paleogenomic perspective on evolution and gene function: new insights from ancient DNA. Science 343.

9. Wilkins D, Lauro FM, Williams TJ, Demaere MZ, Brown MV, et al. (2012) Biogeographic partitioning of Southern Ocean microorganisms revealed by metagenomics. Environ Microbiol 15: 1318–1333.

10. Shapiro BJ, Friedman J, Cordero OX, Preheim SP, Timberlake SC, et al. (2012) Population genomics of early events in the ecological differentiation of bacteria. Science 336: 48–51.

11. Rudi K, Skulberg OM, Jakobsen KS (1998) Evolution of cyanobacteria by exchange of genetic material among phyletically related strains. J Bacteriol 180: 3453–3461.

12. Rae BD, Forster B, Badger MR, Price GD (2011) The CO2-concentrating mechanism of Synechococcus WH5701 is composed of native and horizontally-acquired components. Photosynth Res 109: 59–72.

13. Six C, Thomas JC, Garczarek L, Ostrowski M, Dufresne A, et al. (2007) Diversity and evolution of phycobilisomes in marine Synechococcus spp.: a comparative genomics study. Genome Biol 8.

14. Tung HC, Bramall NE, Price PB (2005) Microbial origin of excess methane in glacial ice and implications for life on Mars. Proc Natl Acad Sci USA 102: 18292–18296.

15. Elena SF, Lenski RE (2003) Evolution experiments with microorganisms: The dynamics and genetic bases of adaptation. Nat Rev Genet 4: 457–469.

16. Lieberman TD, Michel JB, Aingaran M, Potter-Bynoe G, Roux D, et al. (2011) Parallel bacterial evolution within multiple patients identifies candidate pathogenicity genes. Nat Genet 43: 1275–U1148.

17. Denef VJ, Banfield JF (2012) In situ evolutionary rate measurements show ecological success of recently emerged bacterial hybrids. Science 336: 462–466.

18. Barrick JE, Yu DS, Yoon SH, Jeong H, Oh TK, et al. (2009) Genome evolution and adaptation in a long-term experiment with Escherichia coli. Nature 461: 1243–1247.

19. Bramall NE (2007) The remote sensing of microorganisms: University of California, Berkeley.

20. Rohde RA, Price PB, Bay RC, Bramall NE (2008) In situ microbial metabolism as a cause of gas anomalies in ice. Proc Natl Acad Sci USA 105: 8667–8672.

21. French CS, Smith JHC, Virgin HI, Airth RL (1956) Fluorescence-spectrum curves of chlorophylls, pheophytins, phycoerythrins, phycocyanins and hypericin. Plant Physiol 31: 369–374.

22. Giovannoni SJ, Rappe MS, Vergin KL, Adair NL (1996) 16S rRNA genes reveal stratified open ocean bacterioplankton populations related to the green non-sulfur bacteria. Proc Natl Acad Sci USA 93: 7979–7984.

23. Rocap G, Distel DL, Waterbury JB, Chisholm SW (2002) Resolution of Prochlorococcus and Synechococcus ecotypes by using 16S-23S ribosomal DNA internal transcribed spacer sequences. Appl Environ Microb 68: 1180–1191.

24. Willerslev E, Hansen AJ, Poinar HN (2004) Isolation of nucleic acids and cultures from fossil ice and permafrost. Trends Ecol Evol 19: 141–147.

25. Sogin ML, Morrison HG, Huber JA, Mark Welch D, Huse SM, et al. (2006) Microbial diversity in the deep sea and the underexplored “rare biosphere". Proc Natl Acad Sci USA 103: 12115–12120.

26. Altschul SF, Madden TL, Schaffer AA, Zhang JH, Zhang Z, et al. (1997) Gapped BLAST and PSI-BLAST: a new generation of protein database search programs. Nucleic Acids Res 25: 3389–3402.

27. Pruesse E, Quast C, Knittel K, Fuchs BM, Ludwig WG, et al. (2007) SILVA: a comprehensive online resource for quality checked and aligned ribosomal RNA sequence data compatible with ARB. Nucleic Acids Res 35: 7188–7196.

28. Edgar RC (2004) MUSCLE: multiple sequence alignment with high accuracy and high throughput. Nucleic Acids Res 32: 1792–1797.

29. Dethlefsen L, Huse S, Sogin ML, Relman DA (2008) The pervasive effects of an antibiotic on the human gut microbiota, as revealed by deep 16S rRNA sequencing. PLoS Biol 6: 2383–2400.

30. Galand PE, Casamayor EO, Kirchman DL, Lovejoy C (2009) Ecology of the rare microbial biosphere of the Arctic Ocean. Proc Natl Acad Sci USA 106: 22427–22432.

31. Wang Q, Garrity GM, Tiedje JM, Cole JR (2007) Naive Bayesian classifier for rapid assignment of rRNA sequences into the new bacterial taxonomy. Appl Environ Microb 73: 5261–5267.

32. Pruesse E, Peplies J, Glockner FO (2012) SINA: Accurate high-throughput multiple sequence alignment of ribosomal RNA genes. Bioinformatics 28: 1823–1829.

33. Altekar G, Dwarkadas S, Huelsenbeck JP, Ronquist F (2004) Parallel metropolis coupled Markov chain Monte Carlo for Bayesian phylogenetic inference. Bioinformatics 20: 407– 415.

34. Sun SL, Chen J, Li WZ, Altintas I, Lin A, et al. (2011) Community cyberinfrastructure for advanced microbial ecology research and analysis: the CAMERA resource. Nucleic Acids Res 39: D546–D551.

35. Camacho C, Coulouris G, Avagyan V, Ma N, Papadopoulos J, et al. (2009) BLAST+: architecture and applications. BMC Bioinf 10.

36. Lemieux-Dudon B, Blayo E, Petit JR, Waelbroeck C, Svensson A, et al. (2010) Consistent dating for Antarctic and Greenland ice cores. Quat Sci Rev 29: 8–20.

37. Price PB, Woschnagg K, Chirkin D (2000) Age vs depth of glacial ice at South Pole. Geophys Res Lett 27: 2129–2132.

38. Spaulding NE, Higgins JA, Kurbatov AV, Bender ML, Arcone SA, et al. (2013) Climate archives from 90 to 250 ka in horizontal and vertical ice cores from the Allan Hills Blue Ice Area, Antarctica. Quat Res 80: 562–574.

39. Rusch DB, Halpern AL, Sutton G, Heidelberg KB, Williamson S, et al. (2007) The Sorcerer II Global Ocean Sampling expedition: Northwest Atlantic through Eastern Tropical Pacific. PLoS Biology 5: 398–431.

40. Venter JC, Remington K, Heidelberg JF, Halpern AL, Rusch D, et al. (2004) Environmental genome shotgun sequencing of the Sargasso Sea. Science 304: 66–74.

41. DeLong EF, Preston CM, Mincer T, Rich V, Hallam SJ, et al. (2006) Community genomics among stratified microbial assemblages in the ocean’s interior. Science 311: 496–503.

42. Bowler C, Karl DM, Colwell RR (2009) Microbial oceanography in a sea of opportunity. Nature 459: 180–184.

43. Simon C, Wiezer A, Strittmatter AW, Daniel R (2009) Phylogenetic diversity and metabolic potential revealed in a glacier ice metagenome. Appl Environ Microb 75: 7519–7526.

